# Survey of gene, lncRNA and transposon transcription patterns in four mouse organs highlights shared and organ-specific sex-biased regulation

**DOI:** 10.1101/2024.09.10.612032

**Authors:** Qinwei Kim-Wee Zhuang, Klara Bauermeister, Jose Hector Galvez, Najla Alogayil, Enkhjin Batdorj, Fernando Pardo Manuel de Villena, Teruko Taketo, Guillaume Bourque, Anna K. Naumova

**Author notes:** Corresponding authors: Anna K. Naumova, PhD, RI MUHC, 1001 Decarie Blvd., Bloc E, EM03226, Montreal, Quebec, Canada, H4G 1J3, Guillaume Bourque, PhD, McGill Genome Centre, 740 Dr Penfield Ave, Room 6103, Montréal, Québec, Canada, H3A 1A4. These authors contributed equally to the work (Qinwei Kim-Wee Zhuang, Klara Bauermeister).

## Abstract

**Background:** Sex-biased gene regulation is the basis of sexual dimorphism in phenotypes and has been studied across different cell types and different developmental stages. However, sex-biased expression of transposable elements (TEs) that represent nearly half of the mammalian genome and have the potential of influencing genome integrity and regulation, remains underexplored.

**Results:** Here, we report a survey of gene, lncRNA and TE expression in four organs from mice with different combinations of gonadal and genetic sex. Data show remarkable variability among organs with respect to the impact of gonadal sex on transcription with the strongest effects observed in liver. In contrast, the X-chromosome dosage alone had modest influence on sex-biased transcription across different organs, albeit interaction between X-dosage and gonadal sex cannot be ruled out. The presence of the Y chromosome influenced TE, but not gene or lncRNA expression in liver. Notably, 90% of sex-biased TEs (sDETEs) reside in clusters. Moreover, 54% of these clusters overlap or reside close (<100 kb) to sex-biased genes or lncRNAs, share the same sex bias, and also have higher expression levels than sDETE clusters that do not co-localize with other types of sex-biased transcripts. We also tested the heterochromatic sink hypothesis that predicts higher expression of TEs in XX individuals and found no evidence to support it.

**Conclusions:** Our data show that sex-biased expression of TEs varies among organs with highest numbers of sDETEs found in liver following the trends observed for genes and lncRNAs. It is enhanced by proximity to other types of sex-biased transcripts.

## BACKGROUND

Biological sex is an important factor in mammalian biology with sex differences observed in multiple phenotypes from physiology and metabolism to complex behaviors. In humans, male sex is a major risk factor for most types of cancers and poor outcomes following infections, whereas the female sex increases the risk of autoimmune diseases, to give a few examples [1–5]. The major contributors to biological sex differences in mammals are the sex chromosomes (XX in females and XY males) defining the genetic sex, and the gonads defining the gonadal sex ([6, 7], reviewed in [8]). The gonadal sex hormone and growth hormone signaling pathways and networks of their target genes represent the next layers of sexually dimorphic factors that contribute to gene regulation and, hence, phenotypes [9–15]. X-chromosome dosage or the presence/absence of the Y chromosome may impact gene regulation through the actions of X- or Y-linked epigenetic modifiers and transcription factors [16–20].

To understand the regulatory hierarchies governing sex-biased gene expression and get closer to identifying specific signaling pathways or transcription factors, several approaches have been used, including sex-reversed mice [16, 21], the four core genotypes (FCG) mouse model [11], mouse models addressing the roles of pituitary hormones [9, 10], testosterone and estrogen-signaling pathways [11–13], as well as mutations in genes encoding specific transcription factors and epigenetic modifiers [13, 14, 22–26]. Liver is one of the best studied mouse organs with respect to sex-biased gene regulation. Most of the expression data from adult liver show major impacts of gonadal sex, growth hormone, testosterone, and estrogen but minor contributions from the sex chromosomes. In contrast, expression studies of certain mouse and human cell types have demonstrated the significant role of the sex-chromosome complement [16, 26–32]. Such differences likely reflect cell-type specificity of sex-biased regulation. Analyses of sex-biased expression of another group of transcripts, the long non-coding RNAs (lncRNAs) suggest a strong regulatory potential and significant contribution to sex-biased transcriptomes and sexually dimorphic phenotypes [33–37].

In most studies, the focus has been on sex-biased gene or lncRNA expression, whereas the role of biological sex in the regulation of TEs has not been examined in detail. Transposition for the majority of competent Class 1 TEs requires an RNA intermediate and, hence, active transcription [38–41]. During mammalian development, TEs normally become active in the developmental time windows when global epigenome reprogramming takes place: in primordial germ cells (PGCs) and preimplantation embryos. Expression of certain TE families facilitates the activation of the embryonic genome [42, 43]. However, after the blastocyst stage, most TEs are silenced again [44–46]. TEs may be expressed as part of lncRNAs, represent about 40% of their exonic sequences [47] and potentially contribute to their function [48]. It is worth noting that the long terminal repeats (LTRs) of many endogenous retroviruses harbor steroid hormone response elements that mediate activation of transcription in response to hormonal stimuli [49–52].

While most TEs are expected to be transcriptionally repressed in adult somatic cells, their abnormal reactivation may lead to gene dysregulation or increase the mutational burden and the risk of neoplastic transformation ([40, 53–55], reviewed in [56–59]). Indeed, global demethylation of TEs has been reported for certain types of cancers, however the driving forces behind TE activation that could explain sex bias in cancer risk have not been elucidated yet (reviewed in [46]).

Furthermore, the “heterochromatic sink” hypothesis provides an intriguing link between sex chromosomes and TE regulation. It has been proposed that X-chromosome inactivation (XCI), which creates a large demand for heterochromatin, could compete with the rest of the cell genome for silencing factors and causes stochastic upregulation of autosomal genes and TEs in individuals with two or more X chromosomes [60]. This may explain sexually dimorphic phenomena associated with different X-chromosome dosage in mammals, such as increased predisposition to neural tube defects in females [61] or increased variability in the expression of agouti alleles in mice [60]. Conversely, TEs may impact genome-wide transcription by sequestering heterochromatin machinery as demonstrated in *Drosophila*, where the Y chromosome that is gene-poor but TE-rich acts as a “heterochromatic sink” influencing heterochromatin distribution patterns genome-wide and contributing to sex bias in ageing [62].

To the best of our knowledge, the impacts of gonadal vs genetic sex on TE regulation and its relationship with sexually dimorphic gene expression in mammalian somatic cells have not been characterized [26]. Our goal was to examine the sex-biased expression patterns of TEs, the roles of the sex chromosomes and gonadal sex in their regulation and elucidate the relationship between TEs and sex-biased expression of other types of transcripts, i.e. genes and lncRNAs. We conducted a survey of sex-biased expression in four organs from mice with different combinations of gonadal sex and sex-chromosome complement which allows to separate the effects of gonadal sex, Y chromosome and X-dosage. In addition, we used our expression data to test the hypothesis that the inactive X acted as a heterochromatic sink and enhanced autosomal TE or gene expression.

## RESULTS

### Mice with different combinations of gonadal sex and sex-chromosome complement and their genetic backgrounds

To explore the contributions of gonadal sex and sex-chromosome complement in an organ-specific context, we analyzed the transcriptomes of four organs from mice with different combinations of gonadal sex and sex-chromosome complement. To generate such combinations, two mouse strains were used: the B6.Y^TIR^ (Tirano) consomic mice that carry the Y chromosome from a house mouse caught in Tirano, Italy, on a C57BL/6J genetic background [63] and mice carrying the patchy fur (*Paf*) mutation on the X chromosome. About 20% of the offspring of X*^Paf^*Y males are females with one X chromosome (XO.F) [64], whereas about 50 % of B6.Y^TIR^ develop into sex-reversed females with two ovaries (XY.F) [63, 65]. The comparison of sex-reversed XY.F to XY males (XY.M) detects biased transcription driven by the gonadal sex differences, whereas comparing XX female littermates (XX.F) to XY.F detects biased transcription driven by the sex-chromosome complement. To distinguish the impact of X-chromosome dosage from that of the Y chromosome, we compared females with monosomy X (XO.F) to their XX*^Paf^* (XX*^Paf^*.F) littermates originating from crosses between X*^Paf^*Y males and C57BL/6J females.

Since genetic background influences gene regulation, and variability in the genetic background may obscure the effect of sex on expression [66, 67], mice were genotyped using the MiniMUGA version of the Mouse Universal Genotyping Array (MUGA) [68]. Mice from the B6.Y^TIR^ cross carried 100% C57BL/6J alleles on autosomes, the X chromosome and mitochondria, while the Y chromosome was derived from the TIRANO strain. The XO.F and XX*^Paf^*.F mice had varying contributions from the C3H/He strain in heterozygosity (ranging from 0 and 54%) with the remaining alleles derived from the C57BL/6J strain (**Table S1**).

RNA-seq was performed on lung parenchyma, heart, and whole brain samples from adult XX.F, XY.F, XY.M, XO.F, and XX*^Paf^*.F mice. Liver data from our previous study were also analyzed [21]. All data were deposited to NCBI (GSE248074).

### Organ-biased TE expression

As a starting point, we identified *de novo* lncRNAs (referred to as lncRNAs hereafter) in our dataset. In total, 9,754 lncRNAs were identified in brain, 3,823 in heart, 7,885 in liver, and 6,325 in lung (**Table S2**). Data quality check confirmed that all three types of transcripts (genes, lncRNAs, and TE subfamilies) had comparable global expression levels (**Figure S1**) and expression patterns were highly similar between samples from the same organ (**Figure S2**).

Principal component analysis (PCA) for genes, lncRNAs, and TE subfamilies showed clusters formed by organs (**Figure 1A**).

**Figure 1.**
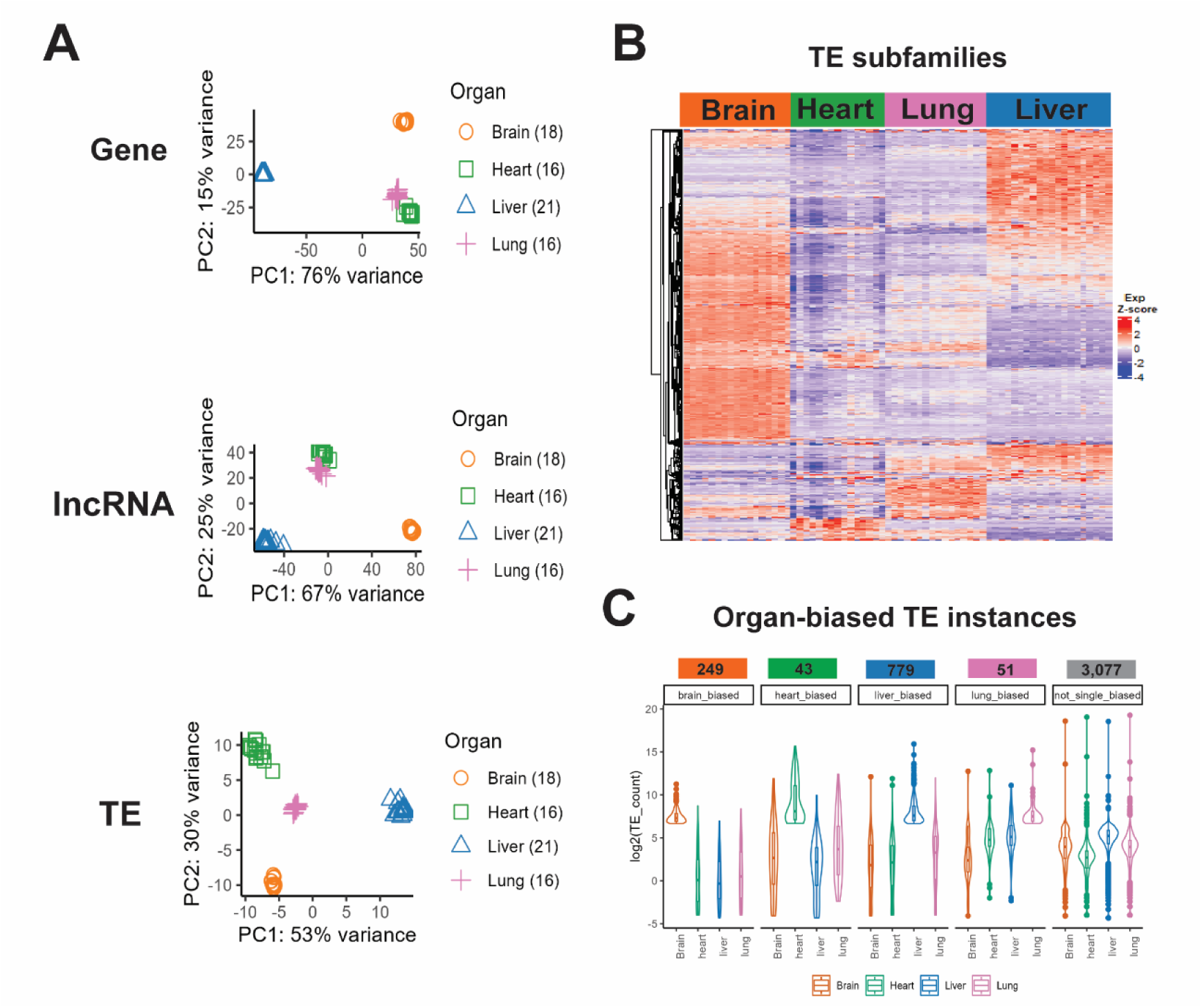
Organ-specific expression patterns. **(A)** PCA plot of gene (top), lncRNA (middle), and TE (bottom) expression. The top 300 genes, lncRNAs or TE subfamilies with the highest variance across samples were used. Numbers in the parenthesis refer to the number of samples for each organ. **(B)** TE subfamily expression profiles show organ-specific patterns. Each column corresponds to one sample and each row - to one TE subfamily (1054 TE subfamilies in total). The Z-score was calculated for each TE subfamily and used to generate the heatmap. **(C)** Organ-biased expression levels of TE instances. Organ-biased TE instances were defined with Z-score > 1, τ > 0.6, and average aligned reads > 100.

To test if organ differences in TE profiles were the result of predominant organ-specific expression of certain TE subfamilies, we identified organ-biased TE subfamilies (with higher expression in one specific organ but lower in the other three) by combining Tao index τ [69, 70] and z-score (τ > 0.6, z-score > 1) (**Figures 1B, S3A, Table S3**). Among the four organ-biased TE subfamilies as well as the subset that was common across organs, the long terminal repeat (LTR) class subfamilies were the most numerous (**Figure S3B**). We further identified organ-biased TEs at the instance level and observed the highest number of organ-biased TE instances in the liver (**Figure 1C, Table S3**).

In summary, our data confirmed organ-specific expression of annotated genes and lncRNAs, in agreement with previous reports [71–73]. In addition, we identified organ-biased TEs at both subfamily and instance levels.

### Gonadal sex and sex-chromosome complement impact mouse transcriptomes in an organ-specific fashion

To explore the impacts of gonadal sex and sex-chromosome complement on expression in different organs, PCA was performed for each type of transcript and each organ, separately (**Figures 2A, S4, S5**). In the gene PCA, brain samples clustered by sex-chromosome complement and genetic background. In the liver, samples formed clusters by both gonadal sex and sex-chromosome complement. It is also notable that in liver, principal component 1 (PC1) had the highest percentage of variance explained (64%) and seemed to be associated with gonadal sex. This suggests that not only were samples in the liver analysis clustering by gonadal sex, but that this was also the variable that explained the greatest difference between samples. In the heart and lung, samples formed clusters based on the presence or absence of the Y chromosome, but the variance explained as estimated by PC1 was lower (29% and 37%, respectively) than in the brain and liver (43% and 64%, respectively).

**Figure 2.**
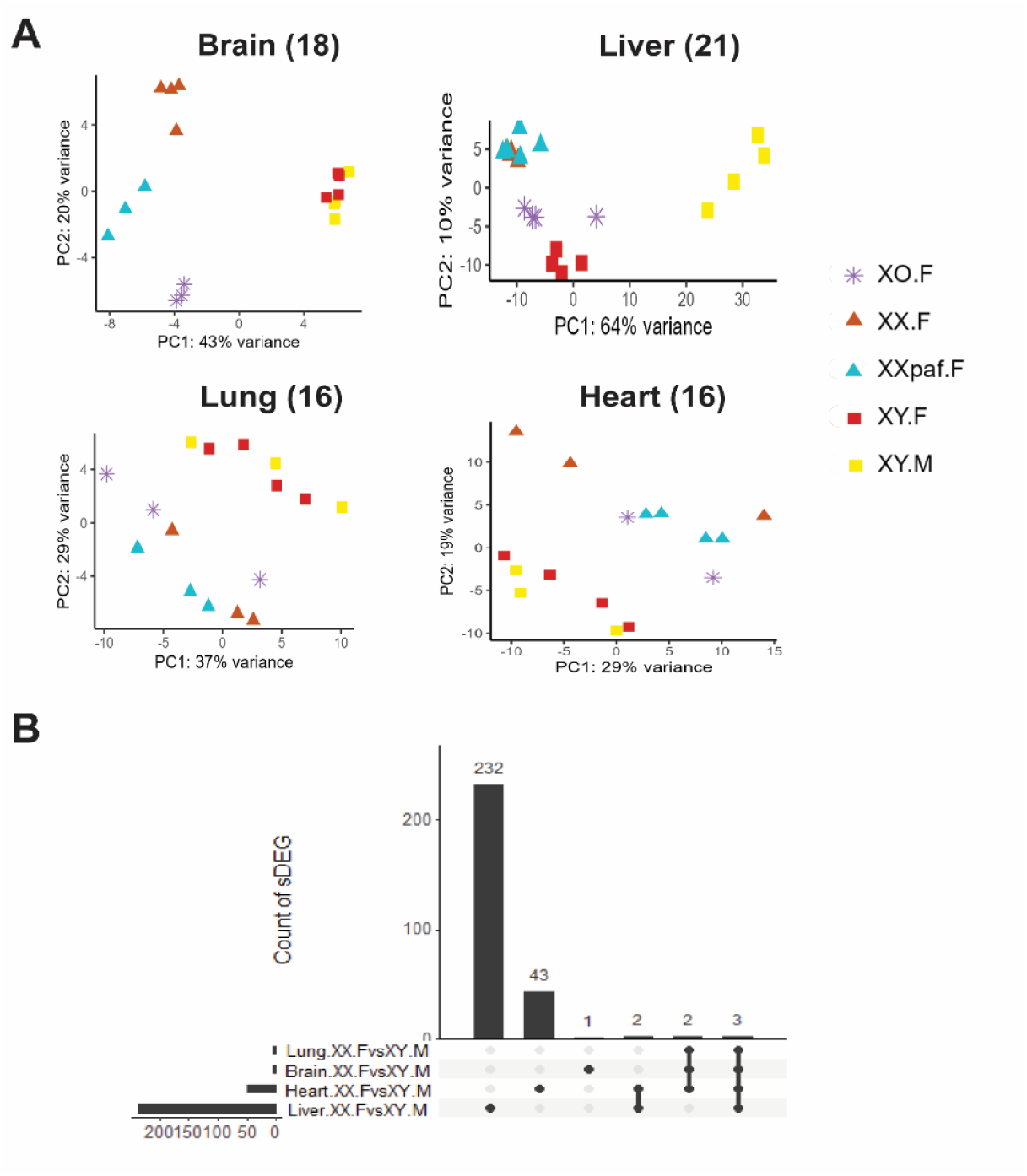
Effects of gonadal sex and sex-chromosome complement on gene expression in four organs. (**A**) PCA plot of gene expression in four organs. (**B**) Numbers of sDEGs from the XX.F vs XY.M comparison in the four organs. Total numbers of DEGs per organ are shown on the bottom left. Numbers of DEGs in each subgroup are shown above each bar.

Separate analysis of X-linked genes showed samples with one and two X chromosomes falling into different clusters, as expected (**Figure S4**). No major effects of gonadal sex or sex-chromosome complement on autosomal gene expression was found in brain, heart or lung, whereas in liver samples formed clusters by gonadal sex (**Figure S4)**. Therefore, the effect of the Y chromosome on the lung and heart transcriptomes was likely driven by Y-linked genes. Similar trends were found for lncRNAs and TEs (**Figure S5**).

Next, we identified sex-associated DEGs (sDEGs) (**Table S4**), lncRNAs (slncRNAs) (**Table S5**), and differentially expressed TEs (sDETEs) (**Table S6**) in four pair-wise comparisons of sex/genotype groups for each organ (numbers summarized in **Table 1**). Overall, the numbers of sDEGs, slncRNAs and sDETEs were concordant within organs with highest numbers of sex-biased transcripts found in the liver. The overlap between sDEGs from different organs was small (**Figure 2B**) with only five DEGs shared between all organs: X-inactive specific transcript (*Xist*) and Y-linked *Kdm5d*, *Uty*, *Ddx3y*, and *Eif2s3y*. Two sDEGs, NADPH oxidase 4 (*Nox4*) and ERBB receptor feedback inhibitor 1 (*Errfi1)*, were found in heart and liver in the XX.F vs XY.M comparisons.

**Table 1.** Numbers of sDEG, slncRNAs, and sDETE in the four organs.

| Comparison | Organ | Lung |  |  | Heart |  |  | Whole brain |  |  | Liver |  |  |
| --- | --- | --- | --- | --- | --- | --- | --- | --- | --- | --- | --- | --- | --- |
|  | Chromosomes | Auto | X | Y | Auto | X | Y | Auto | X | Y | Auto | X | Y |
| XX.F vs XY.M | sDEG | 0 | 1 | 4 | 43 | 3 | 4 | 2 | 1 | 4 | 227 | 6 | 4 |
|  | slncRNA | 2 | 1 | 1 | 1 | 1 | 1 | 0 | 0 | 0 | 207 | 6 | 3 |
|  | sDETE instance | 3 | 0 | 0 | 12 | 0 | 0 | 1 | 0 | 1 | 1401 | 16 | 5 |
| XY.F vs XY.M | sDEG | 6 | 0 | 1 | 32 | 2 | 0 | 0 | 0 | 0 | 174 | 2 | 0 |
|  | slncRNA | 1 | 0 | 0 | 1 | 0 | 0 | 3 | 0 | 0 | 104 | 7 | 0 |
|  | sDETE instance | 2 | 1 | 0 | 9 | 0 | 0 | 0 | 0 | 0 | 855 | 5 | 0 |
| XX.F vs XY.F | sDEG | 0 | 1 | 4 | 5 | 1 | 4 | 0 | 2 | 4 | 3 | 1 | 4 |
|  | slncRNA | 0 | 1 | 1 | 0 | 1 | 1 | 0 | 0 | 0 | 8 | 1 | 3 |
|  | sDETE instance | 1 | 0 | 0 | 3 | 1 | 0 | 1 | 0 | 1 | 147 | 1 | 3 |
| XX<sup>Paf</sup>.F vs XO.F<br>(X-dosage) | sDEG | 2 | 1 | 0 | 1 | 1 | 0 | 3 | 1 | 0 | 1 | 2 | 0 |
|  | slncRNA | 1 | 2 | 0 | 0 | 1 | 1 | 2 | 2 | 0 | 1 | 2 | 1 |
|  | sDETE instance | 1 | 2 | 0 | 0 | 0 | 0 | 5 | 1 | 0 | 16 | 12 | 0 |
| Y-dependent* | sDEG | 0 | 0 | 4 | 1 | 0 | 4 | 0 | 0 | 4 | 3 | 0 | 4 |
|  | slncRNA | 0 | 0 | 1 | 0 | 0 | 1 | 0 | 0 | 0 | 8 | 0 | 3 |
|  | sDETE instance | 1 | 0 | 0 | 3 | 1 | 0 | 1 | 0 | 1 | 146 | 1 | 3 |

sDEG, slncRNA, and sDETE cut-offs: $|\log FC| \geq 1.0$ , adj p-value $< 0.05$ . \* To remove low-expressed transcripts, $\log CPM \geq 1.0$ was applied for sDEGs, and number of average aligned reads $> 50$ was applied for slncRNA and sDETE. Y-dependent transcripts: identified in XX.F vs XY.F but not in XX<sup>Paf</sup>.F vs XO.F.

Overall, gonadal sex had the biggest effect in the liver, followed by a moderate effect in the heart but little effect on brain or lung transcriptomes (**Tables 1, S4, S5, S6**). In contrast to liver where both male- (108 sDEGs, 105 slncRNAs, and 670 sDETEs) and female-biased (121 sDEGs, 96 slncRNAs, and 733 sDETEs) expression was observed, in the heart, all gonadal-sex dependent sDEGs, slncRNA, and most sDETEs (12 out of 13) had higher expression in males.

The numbers of X-dosage dependent and Y-dependent sDEGs were low and similar between organs (**Table 1**), suggesting that the influence of gonadal sex alone or interaction between gonadal sex and sex-chromosome complement rather than sex-chromosome complement alone accounted for most of the organ differences in sex-biased transcription. Y-linked genes *Kdm5d*, *Uty*, *Eif2s3y*, *Ddx3y, Gm29650*, *Gm4017*, and *Gm18665* were expressed in all four organs. Low level expression of *Uba1y, Tspy-ps,* and *Gm28587* was detected in brain only. All these genes are located within a ≈ 0.5 Mb-long contiguous region of Yp, which suggests that this region is transcriptionally active in all four adult organs. A notably high proportion (nearly 10%) of liver sDETEs were sensitive to the presence of the Y chromosome (identified in XX.F vs XY.F but not in XX*^Paf^*.F vs XO.F comparison) (**Table 1**). Among these, 98 had higher expression in XX.F and 52 had higher expression in XY.F samples. The effect of the Y chromosome in brain, heart, and lung was modest compared to that in liver.

In principle, the striking organ differences with respect to sex-biased expression of TEs may be due to increase in the expression of TEs that are sensitive to sex-associated factors in liver and reduced expression of such TEs in the other three organs. To test such a possibility, we compared the overlap between sDETEs and organ-biased TEs. Remarkably, we found only 92 out of 1422 sDETE instances were liver-biased. Hence, liver-biased TEs do not explain the high abundance of sDETE in liver.

We also tested the heterochromatic sink hypothesis, which predicts that genes and TEs would have higher expression in females with two X chromosomes compared to those with one X (e.g. XY.F or XO.F). We did not observe an excess of TEs with higher expression in XX females: 9 of 16 autosomal liver DETEs had higher expression in XX*^paf^*.F compared to XO.F. Hence, our data from adult organs do not support the heterochromatic sink hypothesis.

We next asked if there was interaction between gonadal sex and the sex-chromosome complement. Transcription that requires both factors would be detected only in the XX.F vs XY.M but not XY.F vs XY.M or XX.F vs XY.F comparisons. Thirty-two per cent of liver and 56% of heart sDEGs were unique to the XX.F vs XY.M comparison, suggesting a significant proportion of sDEGs may be regulated by both gonadal sex and sex-chromosome complement. Interaction was observed for slncRNAs and sDETEs too, where 51% of slncRNAs and 49% of sDETEs were unique to the XX.F vs XY.M comparison.

In conclusion, the impact of gonadal sex on expression was wider and most pronounced in the liver and heart, whereas the lung and whole brain showed very modest or no effect of gonadal sex. The effects of the sex-chromosome complement on autosomal expression were limited to a small number of transcripts in all four organs. Yet, the interaction between X-dosage and gonadal sex cannot be ruled out.

### Both gonadal sex and sex-chromosome complement influence expression of X-linked genes, lncRNAs and TEs

To examine the contributions of gonadal sex and sex-chromosome complement on X chromosomal transcription, we next analyzed expression of X-linked genes, lncRNAs and TEs. X-linked sDEGs, slncRNAs and sDETEs were selected using relaxed criteria of adj deseq2 p-value <0.1, either logCPM>1.0 (for sDEG) or baseMean > 50 (for slncRNAs and sDETEs), no cut-off for logFC (**Tables 2, S7**). This relaxed cut-off was used because genes that escape X-inactivation do not necessarily show double the expression in individuals with two X chromosomes compared to those with one [74, 75]. The XX*^Paf^*.F vs XO.F comparison revealed the impact of X-dosage, whereas sex-biased transcripts present in the XX.FT vs XY.FT but not in the XX^Paf^.F vs XO.F comparison were considered to be Y–chromosome dependent (**Table 2**).

**Table 2.** X-linked sDEGs, slncRNAs, and sDETEs.

| Comparison | Transcript type | Lung | Heart | Whole brain | Liver |
| --- | --- | --- | --- | --- | --- |
| XX.F vs XY.M | sDEG | 7 | 6 | 7 | 15 |
|  | slncRNA | 1 | 1 | 0 | 13 |
|  | sDETE instance | 0 | 1 | 0 | 59 |
| XY.F vs XY.M | sDEG | 1 | 9 | 0 | 10 |
|  | slncRNA | 0 | 0 | 0 | 7 |
|  | sDETE instance | 1 | 0 | 0 | 50 |
| XX.F vs XY.F | sDEG | 10 | 4 | 7 | 2 |
|  | slncRNA | 1 | 1 | 0 | 4 |
|  | sDETE instance | 0 | 1 | 0 | 18 |
| XX<sup>Paf</sup>.F vs XO.F (X-dosage) | sDEG | 8 | 2 | 5 | 2 |
|  | slncRNA | 1 | 1 | 3 | 5 |
|  | sDETE instance | 2 | 0 | 1 | 15 |
| Y-dependent* | sDEG | 2 | 2 | 2 | 1 |
|  | slncRNA | 0 | 0 | 0 | 3 |
|  | sDETE instance | 0 | 1 | 0 | 16 |

Relaxed cut-off: adj deseq2 p-value <0.1, either logCPM>1.0 (for sDEG) or baseMean > 50 (for slncRNAs and sDETEs), no cut-off for logFC
\* Y-dependent transcripts: identified in XX.F vs XY.F but not in XX<sup>Paf</sup>.F vs XO.F.

In the brain, all X-linked sDEGs depended on the sex-chromosome complement. All of them, except for glycoprotein m6b (*Gpm6b*) and dystrophin, muscular dystrophy (*Dmd*), had higher expression in females with two X chromosomes, which may be due to escape from X-inactivation (**Table S7**). In lungs, a relatively small number of X-linked sDEGs was detected. Six out of the 7 X-linked sDEGs in the XX.F vs XY.M comparison and all those detected in XX.F vs XY.F or XX*^Paf^*.F vs XO.F comparisons had higher expression in XX females. In the liver and heart, influence of gonadal sex on expression of X-linked genes was evident, which follows the trends observed for autosomal sDEGs. The impact of X-dosage was also detected (**Table 2**). Among genes known to escape X-inactivation, two, *Kdm5c* and *Ddx3x,* had significantly higher expression in the XX compared to XY or XO lungs and brains, but not hearts and livers, whereas *Kdm6a* and *Eif2s3x* displayed consistently higher expression in XX females across all organs. (**Table S7**).

One gene, *Hccs* showed biased expression in three organs but only in the XO.F vs XX*^Paf^*.F comparison with higher expression in XX*^Paf^*.F. *Hccs* is located within the region of heterozygosity where XX*^Paf^* females carry C3H/HeNRj alleles on their paternal X chromosomes. Therefore, it is possible that the *Hccs* expression bias reflects allelic effects rather than escape from XCI.

Similar to autosomal sDEGs, the largest number of X-linked DETEs and slncRNAs was detected in liver with well-pronounced effects of gonadal sex and X-chromosome dosage. Only a few X-linked sDETEs/slncRNAs were found in brain, heart, and lung, even with a relaxed cut- off (**Table 2**). There was no overlap between X-linked sDETEs from liver and those from other organs. Moreover, the Lx8b_dup13983:Lx8b:L1:LINE (genomic position chrX:169360167- 169361279), which was the only X-linked sDETE identified in the XX*^Paf^*.F vs XO.F comparison in both brain and lung, resides distal to *Hccs* and likely reflects the genetic make-up of the X*^Paf^* chromosome.

Thus, expression of X-linked genes, lncRNAs and TEs is regulated by X-dosage in all organs and gonadal sex in liver and heart, consistent with the trends observed for autosomal transcripts.

### Concordant expression between sDEG/slncRNA and sDETEs

Concordance in the numbers of sDEGs, slncRNAs and sDETEs across organs led us to hypothesize that the three types of transcripts were co-regulated. If this was the case, co-regulated transcripts would be found in closer proximity than those that are not associated by regulation. To test this hypothesis, we focused on the liver transcriptome, for which we had the largest number of sDETEs. First, we mapped sDEGs, slncRNAs and sDETEs across chromosomes and confirmed that they were distributed across all chromosomes rather than located on a few specific chromosomes (**Figure S6**). Next, we tested co-localization of sex-biased transcripts within and between the three groups (sDEGs, slncRNAs, sDETEs) using data from the liver XX.F vs XY.M and XY.F vs XY.M comparisons (**Figure 3A, B-C left panels**). About 90% of sDETEs, 30% of sDEGs, and 35% of slncRNAs were located within 100 kb of another sDETE, sDEG or slncRNA, respectively (**Figure 3A**). Moreover, about 55% of sDETEs were located within 100 kb of their nearest sDEG (**Figure 3B**) and about 35% of sDETEs were located within 100 kb of their nearest slncRNA (**Figure 3C**). Hence, most sDETEs tend to co- localize with each other and/or another sex-biased transcript. Therefore, we merged neighboring sDETEs (distance < 100 kb) with the same sex bias into sDETE clusters using data from the XX.F vs XY.M comparison and examined the co-localization between clusters of sDETEs and sDEGs or slncRNAs. Ninety one percent of sDETEs (1296 out of 1,422) formed 173 (101 female-biased vs 72 male-biased) clusters with an average of 7.5 sDETEs per cluster (range 2 to 49) (**Table S8**). In total, 45.1% (78 out of 173) of sDETE clusters overlapped with sDEGs and/or slncRNAs (46 with sDEGs only, 20 with slncRNAs only, and 12 with both sDEGs and slncRNAs). The percentage increased to 54% (94 out of 173) when expanding the distance between the sDETE and nearest sDEG/slncRNA to 100 kb. Hence, slightly more than half of sDETE clusters co-localized with sDEGs or slncRNAs. The rest did not have sex-biased transcripts of either type within a 100 kb distance. Ninety-five percent of sDEG-overlapping and 91% of slncRNA-overlapping sDETE clusters had the same sex bias as the overlapped sDEG (**Figure 3B**) or slncRNA (**Figure 3C**), respectively. The proportion of same sex bias remained above 85% when extending the distance between the sDETE cluster and other sex-biased transcripts to 100 kb but dropped to around 50% when the distance was larger than 100 kb (**Figure 3B-C**). When direction of transcription was examined, 95% (55 out of 58) and 91% (29 out of 32) of sDETE clusters were transcribed from the same strand as the overlapping sDEG or slncRNA, respectively. Examples of sDETE clusters that co-localize with sDEG/slncRNA are shown in **Figure 3D**. Worth noting, singleton sDETEs (located outside of sDETE clusters) showed significantly lower expression levels than those forming clusters, especially those clusters that overlapped with sDEGs (**Figure 3E**).

**Figure 3.**
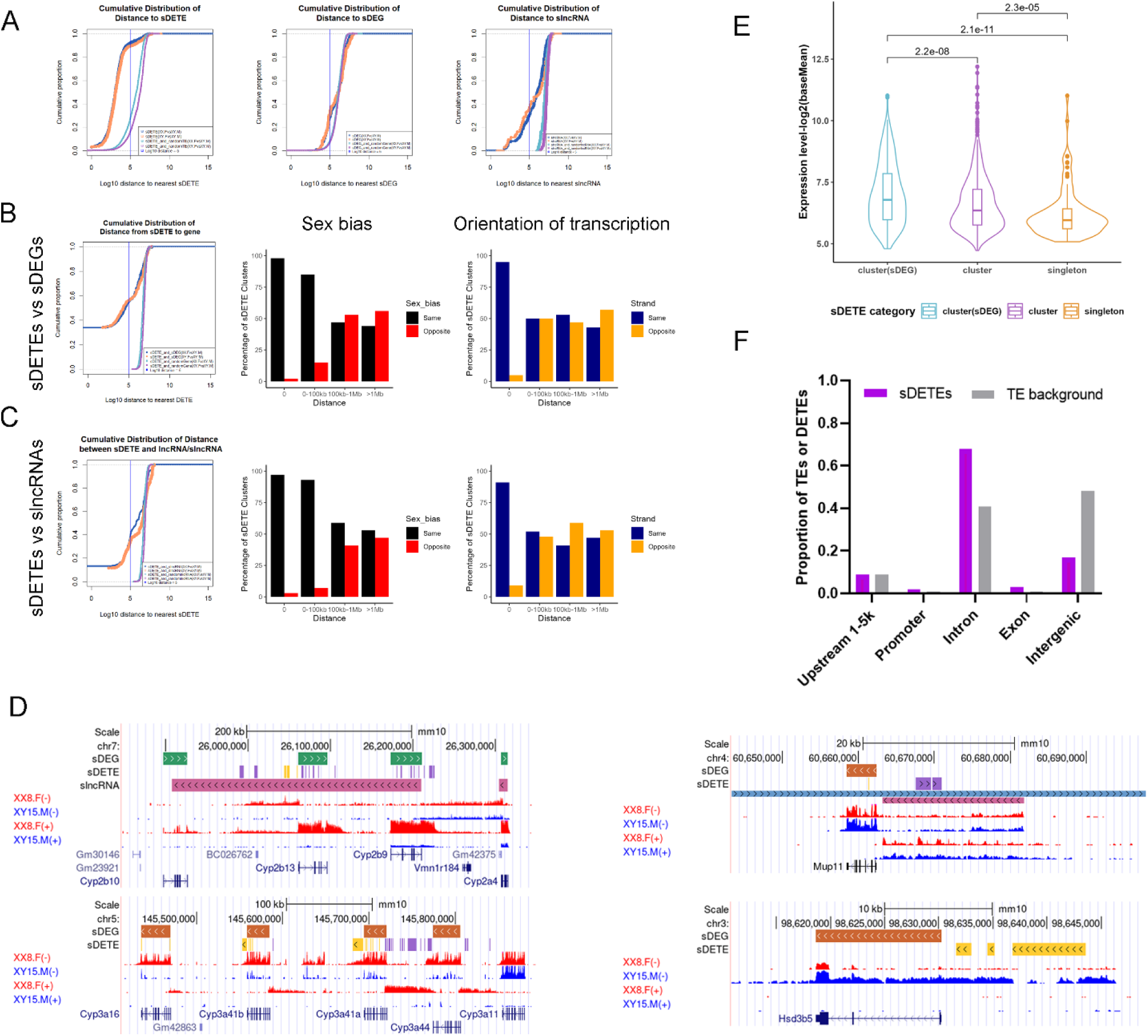
Concordant expression between sDETE and nearest sDEG or slncRNA. (**A**) Cumulative distribution of distances between sDETE and the nearest sDETE (left panel), sDEG and the nearest sDEG (central panel), and slncRNA and nearest slncRNA (right panel). The blue vertical line refers to the distance of 100 kb. (**B**) Concordant expression of sDETEs/sDETE clusters and nearby sDEGs. Cumulative distribution of distance between a sDETE and the nearest sDEG (left panel); sex bias (center) and orientation of expression (right panel) in sDETE clusters and the nearest sDEG as a function of distance. (**C**) Concordant expression of sDETE clusters and nearby slncRNAs, similar to (**B**). (**D**) Examples of co-localization of sDETE clusters, sDEGs and slncRNAs. Examples of female-biased transcripts are shown in the two left panels, examples of male-biased transcripts are shown on the right. Colors and arrows indicate direction of transcription. For sDEGs, green shows the **+** strand, orange indicates the − strand. For sDETEs: purple (+) and yellow (−) strands, respectively. For slncRNAs: blue (+) and pink (−) strands. All features are shown in the context of the UCSC browser. (**E**) Expression levels of singleton sDETEs vs sDETEs residing in clusters. Each dot refers to one sDETE. Three groups were plotted: sDETE that belong to clusters that overlap with sDEGs, sDETEs that belong to clusters but do not overlap with sDEGs, and sDETE singletons. (**F**) Genic annotations of sDETEs from the liver XX.F vs XY.M comparison vs all TEs.

TEs are often part of non-coding transcripts, and 80% of lncRNAs harbor TEs within their exons [47]. To determine whether the sDETEs that overlapped sDEGs or slncRNAs were parts of their mature transcripts, we inspected the genic annotation of sDETEs from the liver XX.F vs XY.M comparison. Briefly, while 33.5% (476 out of 1,422) of sDETE overlapped with sDEGs, only 3.6% (17) were exonic. Meanwhile, 12.8% (182 out of 1,422) of sDETEs overlapped with slncRNAs, of which 62.6% (114) were exonic. In general, TEs tend to reside in intronic and intergenic regions, however, sDETEs were enriched in introns of overlapping coding and non-coding transcripts, but less so in intergenic regions (**Figure 3F**). A possible explanation of such enrichment is that intronic sDETEs are parts of pre-mRNA molecules or intronic debris captured by the RNA-sequencing. If this were the case, one would expect that all intronic TE sequences would be equally represented among transcripts. To test this possibility, we compared the expression levels of sDEG-overlapping TEs based on their sex bias. sDETEs that overlapped with sDEGs had higher expression levels than other TEs overlapping the same sDEG (**Figure S7**). Hence, sDETEs are not a technical artefact but *bona fide* transcripts.

In conclusion, sDETEs tended to form clusters and showed high concordance with overlapping sDEG/slncRNA. sDETE clusters located within 100 kb also showed highly concordant sex bias with neighboring sDEG/slncRNA, regardless of the strand. This suggests possible shared regulation or an impact of domains with active chromatin on TE transcription. However, nearly 50% of sDETE clusters do not colocalize with either type of the sex-biased transcripts and therefore are independently regulated. These clusters have lower transcription levels.

## DISCUSSION

### The contributions of sex-chromosome complement and gonadal sex to sex-biased gene, lncRNA and TE regulation vary in different organs

Here, we conducted a survey of sex-biased gene, lncRNA and TE expression and compared the contributions of the sex-chromosome complement and gonadal sex in four mouse organs: lung parenchyma, heart, whole brain, and liver. For all three types of transcripts, the impact of gonadal sex was wider and most pronounced in liver and heart, whereas the lung and whole brain transcriptomes showed very modest or no effect of gonadal sex. Our data support previously reported dominant roles of gonadal-sex associated factors, including those from growth hormone signaling pathways, on gene expression in liver [9–13]and demonstrate the influence of gonadal sex on TE expression. They also suggest that the four organ differences in sex-biased expression may be largely driven by different sensitivity to factors associated with gonadal sex rather than sex-chromosome complement. The overall low numbers of sDEGs in lung and brain are consistent with those observed in other studies of the C57BL/6J strain [76–78]. However, the number of heart sDEGs found in our study is considerably lower than those reported by others [27]. Such differences may be associated with the use of more stringent inclusion/exclusion criteria. In addition, the slncRNAs that we identified among the *de novo* lncRNAs provided novel targets, beyond annotated genes, of sex-biased transcripts for further investigation. Overall, we observed correlation between numbers of sex-biased genes, lncRNAs and TEs among organs, which suggests common tissue-specific regulatory mechanisms affecting sex-biased transcription.

The effects of the sex-chromosome complement on autosomal gene expression were limited, mostly to four Y-linked genes that were expressed across all tested organs and a few autosomal genes, which showed organ-specific sex-biased expression. In the mouse liver, autosomal TEs but not genes or lncRNAs were sensitive to the presence of the Y chromosome. This is consistent with our previous finding of the influence of the Y chromosome on methylation levels of autosomal repetitive elements [79].

When expression of X-linked transcripts was examined using relaxed inclusion criteria, we found a relatively small number of X-linked DEGs,slncRNAs and DETEs. In the brain, heart and lung, most of X-linked transcripts were sensitive to X-dosage. Most of the X-dosage sensitive DEGs are either known or were predicted to escape X-inactivation [75, 80, 81]. Among them are the well-known escapees *Kdm6a*, *Kdm5c*, *Ddx3x*, and *Eif2s3x.* However, in the liver, gonadal sex had a stronger influence on X-linked transcription, which is consistent with the patterns observed for autosomal transcripts. Our data suggest that transcriptional regulation of X-linked genes and TEs depends on general regulatory mechanisms shared with the rest of the genome as well as the X-inactivation status and these impacts are organ-specific.

### Sex-biased TE expression is organ-specific and associated with sex-biased gene expression

We observed variation in the repertoires and numbers of expressed TEs at both subfamily and instance levels among the four tested organs. Liver had the largest number of highly expressed TEs compared to the other three organs investigated. Organ-biased TE expression spans multiple TE classes, however the LTR class was the most enriched among organ-biased TE subfamilies as well as subfamilies that were common between organs, suggesting that LTRs are most actively transcribed TEs in all four mouse organs.

Organ differences in TE expression were also observed when comparing between sex/genotype groups within each organ separately. Liver had by far more sDETE subfamilies and instances than brain, heart or lung. On autosomes, both gonadal sex and sex-chromosome complement (mainly the presence of the Y chromosome) influenced sex-biased expression of TEs in liver, but the impact of gonadal sex was more observable than that of the sex-chromosome complement. However, there were sex-biased transcripts that were found only in the XX.F vs XY.M comparison suggesting potential interaction between gonadal sex and sex-chromosome complement. In conclusion, the impact of gonadal sex and sex-chromosome completement on TE expression is organ-specific, and most observable in the liver. This warrants further investigation of the tissue-specific mechanisms involved.

The similarities between gene or lncRNA and TE expression patterns with respect to sex and organ-specificity led us to examine the relationship between the three types of sex-biased transcripts in further detail. We found clustering of sDETEs and concordant regulation of sex-biased expression of TEs and genes/lncRNAs in the adult mouse liver, which is consistent with findings in other species [82, 83]. We speculate that sex-biased expression of other types of transcripts boosts expression of neighboring TEs in two different ways: sDETEs are activated by expression of overlapping sDEGs or slncRNAs, whereas sDETEs located within 100 kb of the nearest sDEG or slncRNA are likely the result of local active chromatin environment. The effect fades with increasing distance from the sDEG. We also speculate that such a concordance between gene, lncRNA and TE expression may have implications for TE-driven mutagenesis in somatic cells of an adult organism.

### The heterochromatic sink hypothesis

The heterochromatic sink hypothesis predicts that XX animals should have higher expression levels of autosomal genes or TEs compared to animals with only one X [60, 84]. In our liver dataset, we found only 2 X-dosage dependent autosomal DEGs and 16 sDETEs. X-dosage dependent autosomal DETEs show comparable proportions of XO-biased and XX-biased expression. Overall, the small numbers of X-dosage sensitive DEGs and DETEs do not support a major genome-wide effect of X-dosage on autosomal gene regulation in either of the four adult organs. Hence, if the inactive X acts as a heterochromatic sink activating expression across autosomes, its effect is not detected in adult somatic cells using our present approaches. We speculate that in mammals, the phenomenon is either limited to early developmental stages [85], or increases the rate of stochastic epigenetic anomalies, or absent in normal conditions and requires mutations or environmental insults that would destabilize the epigenetic silencing machinery to exert its regulatory impact. However, we observe a rather high number of DETEs whose expression seems to be sensitive to the presence/absence of the Y chromosome. The mouse Y chromosome has a unique structure with at least 200 copies of the same DNA region and is also enriched for TEs [86]. Our observation warrants further experiments to test whether the Y chromosome may act as a heterochromatic sink in mouse cells.

In conclusion, our results support previous findings that sex-biased expression varies among organs. They also suggest that gonadal sex may be the driving force for organ differences in sex biased expression. It has the most pronounced impact on the adult liver and heart transcriptomes, whereas lung parenchyma and whole brain have few sex-biased genes and TEs. We find a significant impact of sex on TE transcription. Importantly, sDETEs tend to reside in clusters with about 50% of these clusters colocalizing with other sex-biased transcripts sharing the same sex-bias.

## METHODS

### Mice and crosses

**C57BL/6J** mice were purchased from the Jackson Laboratory (Bar Harbor, Maine, USA).

**B6.Y^TIR^** consomic mice were maintained in our colony (TT) by breeding of B6.Y^TIR^ males to C57BL/6J females. B6.Y^TIR^ males were crossed to wild-type C57BL/6J females to generate XY^TIR^ sex-reversed females (XY.F) and males (XY.M), as well as XX females (XX.F). Female offspring were genotyped using PCR amplification of the zinc finger protein on the Y (*Zfy*) sequence of DNA from ear punches, using the primers and conditions described in [87].

Liver (4 XX.F, 4 XY.M, and 4XY.F), lung parenchyma (3 XX.F, 3 XY.M, and 4XY.F), heart (3 XX.F, 3 XY.M, and 4XY.F), and whole brain (4 XX.F, 4 XY.M, and 4XY.F) were collected from 8-week old mice and used for RNA extraction. Gonadal sex was confirmed at the time of organ collection.

**B6.C3H/HeSn-Paf** mice (referred to as *Paf* from this point on) were generated by backcrossing C3H/HeSn-Paf/J carriers of the patchy fur (*Paf*) mutation purchased from the Jackson Laboratory to C57BL/6J mice (Jackson Laboratory, Bar Harbor, Maine, USA). Males that carry the *Paf* mutation were identified based on their hair loss phenotype and crossed to C57BL/6J females. Female offspring from these crosses were genotyped using RT-PCR for the *Xist* gene, which is expressed in XX females (XX*^Paf^*.F) but not in XO females (XO.F) [88]. Liver (5 XX*^Paf^*.F and 4 XO.F), lung parenchyma (3 XX*^Paf^*.F and 3 XO.F), heart (4 XX*^Paf^*.F and 3 XO.F), whole brain (3 XX*^Paf^*.F and 3 XO.F) samples from 8-week old N6 and N7 XO females (XO.F) and their XX*^Paf^* (XX*^Paf^*.F) littermates were collected and used for RNA extraction.

To reduce variability that could arise from circadian rhythmicity, samples were collected always on the same time of the day between Zeitgeber time (ZT) 6 and ZT8.

All procedures were conducted in accordance with the guidelines set by the Canadian Council on Animal Care (Ottawa, Ontario, Canada) and were approved by the Animal Care Committee of the McGill University Health Center (Montreal, Quebec, Canada).

### Genotyping

DNA was extracted from mouse tissues using the DNeasy Blood & Tissue Kit (catalog # 69596, QIAGEN, Valencia, CA) and shipped to Neogen Inc (Lincoln, NE) for genotyping using the MiniMUGA array [68]. Chromosomal sex was determined as originally described [68]. Note that MiniMUGA can accurately discriminate between XX, XY, XO and XXY complements independently of genetic background. The contribution of different inbred strains to the genetic make-up of each sample was estimated as the percent of the autosomal and X chromosome genome using exclusively markers that were informative for these backgrounds. IBD, or identity by descent represents regions of the genome where MiniMUGA is unable to robustly discriminate between the relevant genetic backgrounds.

### RNA Extraction and Sequencing

Total RNA was extracted from mouse organs using Trizol Reagent (Thermo Fisher Scientific, MA, US) and purified using the RNeasy MinElute Cleanup Kit (Qiagen, NL).

Library preparation and RNA-sequencing (RNA-seq) were performed at the McGill University and Genome Quebec Innovation Centre for the liver, and at the McGill Genome Centre for the brain, lung, and heart tissues. Paired-end sequencing was performed using a NovaSeq6000 S4 sequencer. For liver comparisons, previously generated RNA-seq data were used [21].

### Data preprocessing

Data preprocessing and alignment was performed using the GenPipes RNA-seq pipeline [89], with default parameters applied unless mentioned otherwise. Reads were trimmed and filtered for quality, then aligned to the mouse reference genome (GRCm38) using STAR [90], and transcript abundance was estimated using HT-Seq Count (version 0.11.0 for brain, 0.6.0 for liver) [91] for gene expression. We applied the default setting of STAR [90] to retrieve only properly paired alignments and both parameters --winAnchorMultimapNmax and – outAnchorMultimapNmax were set as 100 for the optimization of assigning multi-mapped reads. Alignment BAM files were input to package TEtranscripts (version 2.0.3) [92] to obtain read counts at a TE subfamily level. TE annotation files were retrieved from (https://labshare.cshl.edu/shares/mhammelllab/www-data/TEtranscripts/TE_GTF/mm10_rmsk_TE.gtf.gz). The raw count of TE subfamily was defined as the global sum of reads aligned to all TE instances of each subfamily. Options [-b] and [--sortByPos] were applied to run *TEcount* in TEtranscripts package. Default parameters were applied if not mentioned otherwise. To obtain TE read counts at a locus/instance level, alignment BAM files were input to package TElocal (version 1.1.1) (https://github.com/mhammell-laboratory/TElocal). Options [-b], [--sortByPos], and default number of iterations [--iteration 100] that optimized multi-reads assignment were applied. Genes and TEs with extremely low read counts (average < 5 in all samples) were excluded to reduce noise.

### Identification of de novo lncRNAs

To identify *de novo* lncRNAs in our dataset, output alignment bam files from STAR [90] were merged for samples from the same organ, followed by transcript calling with Stringtie (version 2.1.4) [93] to assemble and extract transcripts with length > 200 bp. Next, the list of transcripts was input into FEElnc, a computational tool used to predict lncRNAs [94]. FEElnc identified novel transcripts that did not overlap with annotated genes of the reference genome (GRCm38.Ensembl102.gtf) [95],then predicted coding potentials of input transcripts with a Random Forest model trained with intrinsic properties of user-input mRNA sequence (such as mRNA sizes, k-mer frequencies, and ORF coverage). Output files of FEElnc included lists of predicted lncRNAs and mRNAs. We curated the list of lncRNA to include the ones with a minimum of 50 aligned reads as the estimated *de novo* lncRNA for our dataset.

### PCA and heatmap

Principal component analysis (PCA) was performed for the top 300 genes with highest variance of expression level, for top 300 lncRNA, and for top 300 most TE subfamilies or TE instances. Heatmap was generated with ComplexHeatmap (v3.10[96]). To visualize relative expression levels of TE subfamilies across organs, organ-relative expression z-score was calculated. To calculate the z-score, mean and squared root (sqrt) of variance were firstly calculated for each TE subfamily across four organs. Z-score = (expression of TE subfamily – mean) / sqrt(variance) was calculated for each TE subfamily.

### Organ-biased TE analysis

To identify TE subfamily with higher expression in one specific organ but lower in all other three organs, we applied organ-relative expression z-score (z-score > 1) and Tao Index (τ> 0.6, [69]. Z-score > 1 was used to select TE subfamilies with relatively higher expression among the four organs, while τ> 0.6 was applied to underscore high expression in only one organ. The Tao Index τ threshold was tuned to best reflect organ differences with both good sensitivity and specificity (**Figure S8**). To calculate τ, mean value µ was calculated in each organ for each TE subfamily. The maximum value (*Max*) across four organs was also calculated for each TE subfamily and the Tao Index τ for each TE subfamily was then calculated with the following formula (modified from [69]). An additional parameter (average expression level across all four organs: baseMean > 500) was also applied to inspect the subset of highly expressed organ-biased TE subfamilies. Organ-biased TE instances within such highly expressed organ-biased TE subfamilies were also identified with parameters z-score > 1 and τ> 0.6 and baseMean > 100. We used baseMean > 100 (instead of baseMean >500) because individual TE instances showed overall lower aligned reads than all instances combined for the TE subfamily level.

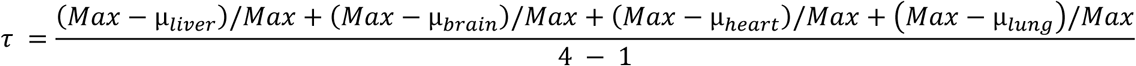

### Differential analysis

For all four investigated organs, sDEGs, slncRNAs, and sDETEs were identified using the DESeq2 (v.1.20.0) [97]and EdgeR (version 3.22.5) ([98]) packages were used. Cut-off threshold for sDEGs was set as |logFC|≥1.0, logCPM≥1.0, adjusted p-value <0.05. Only lncRNAs and DETEs with a number of aligned reads > 50 were kept in the preprocessing step. Cut-off thresholds for both were set as |logFC|≥1.0, adjusted p-value <0.05.

### Genic annotation of sDETE instances

The genic annotation was performed using R package GenomicFeatures (v1.38.2, [99]). The genic annotations included 1–5 kb upstream of the transcription start site (Upstream 1–5 kb), the promoter (<1 kb upstream of TSS), introns, and exons.

#### Visualization of location of sDETE instances

Visualization of chromosomal location of sDETE, sDEG, and slncRNAwas performed with R package chromoMap (version 4.1.1) [100]).

#### Cumulative distribution of distances between closest sex-biased signals

Distance between sex-biased signal and the nearest peer or signal from another sub-group was calculated using the bedtools closest (v2.29.2, [101]).

Cumulative distribution of distance between sDETEs/sDEG/slncRNAs and the nearest sDETE/sDEG/slncRNA was calculated for the liver comparison XX.F vs XY.M.ZTo generate a random background group for sDEGs, we randomly selected 200 genes, 2,000 TEs, or 2,000 lncRNAs and calculated the distance between the selected genes to nearest sDEG/sDETE/slncRNA. The generation process was repeated 10,000 times to build a background group.

#### Cluster identification

Clusters were identified for liver sDETEs for the comparison XX.F vs XY.M. Nearby sDETEs (two or more) were merged into a sDETE cluster if their distance to each other was < 100 kb. The cutoff threshold of 100 kb was chosen based on the cumulative distribution of distance between nearby sex-biased signals (as shown in **Figure 3A).**

## DATA ACCESS

All data are deposited to NCBI GEO database (GSE248074).

Scripts and pre-processed files to generate figures for the paper are uploaded on GitHub (https://github.com/qzhuang8/sex-biased-TEs-paper-script).

## ACKNOWLEDGEMENTS

The study was supported by funds from the Natural Sciences and Engineering Research Council of Canada Discovery Grant program and Discovery Accelerator Supplement (to AKN) and in part by a Canada Institute of Health Research (CIHR) program grant (CEE-151618) for the McGill Epigenomics Mapping Center, which is part of the Canadian Epigenetics, Environment and Health Research Consortium (CEEHRC) Network (to GB). G.B. is supported by a Canada Research Chair Tier 1 award and a FRQ-S Distinguished Research Scholar award.

QKWZ is a recipient of Doctoral Training Scholarship from Fonds de recherche du Québec (FRQS) and Japan Science and Technology Agency Support for Pioneering Research Initiated by the Next Generation (JST SPRING). NA is a recipient of the RI MUHC graduate scholarship. This research was enabled in part by support provided by Calcul Quebec (calculquebec.ca) and the Digital Research Alliance of Canada (alliancecan.ca).

We are grateful for Pablo Hock, Timothy A Bell and Matt Blanchard for help processing DNA samples for genotyping.

