## Supplemental Figures for "Survey of gene, lncRNA and transposon transcription patterns in four mouse organs highlights shared and organ-specific sex-biased regulation"

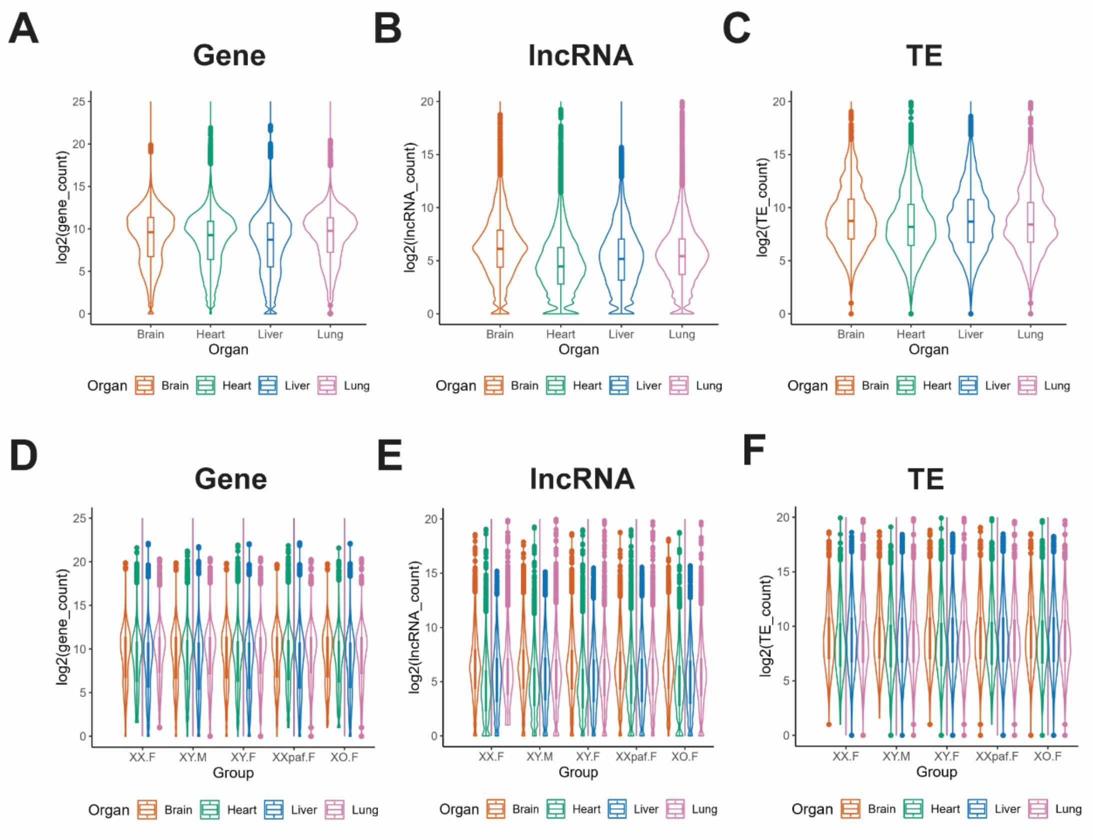


**Figure S1. Quality check confirms comparable transcript abundance for genes, *de novo* lncRNAs and TE subfamilies across organs.** (**A-C**) Expression levels of genes, (*de novo*) lncRNAs, and TEs by organ, with all sex/genotype groups. (**D-E**) Similar to A-C but with different sex/genotype groups being plotted separately.

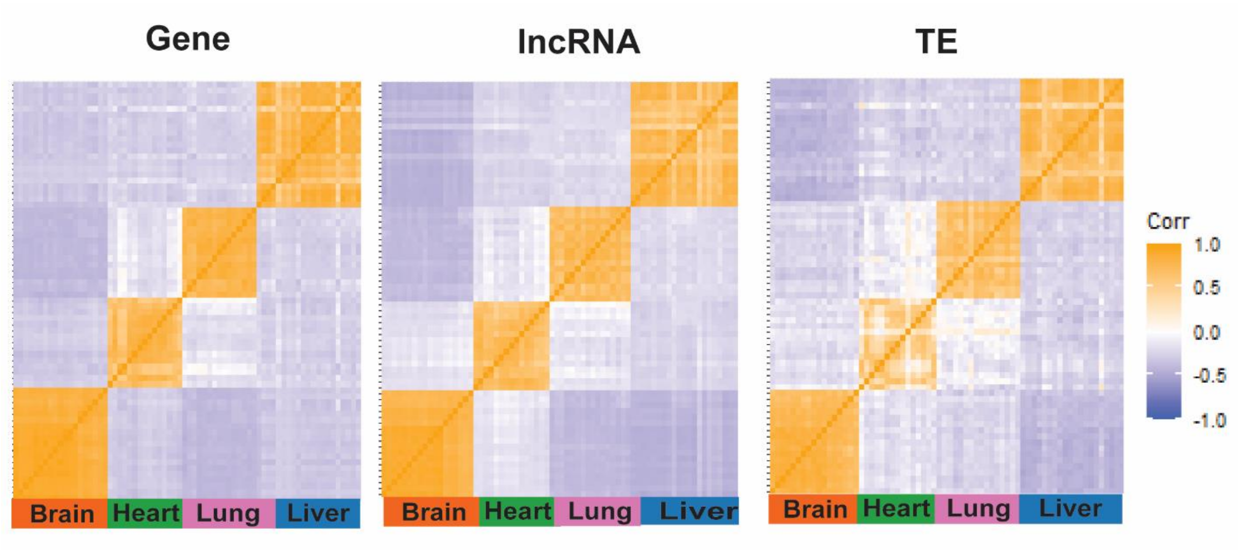


**Figure S2. Heatmaps highlight organ-specific expression patterns for genes, (*de novo*) lncRNAs, and TE subfamilies.** Heatmap of correlation coefficients of expression levels between samples. Both rows and columns refer to from the four invested organs. Correlation coefficients are calculated between pairs of samples.

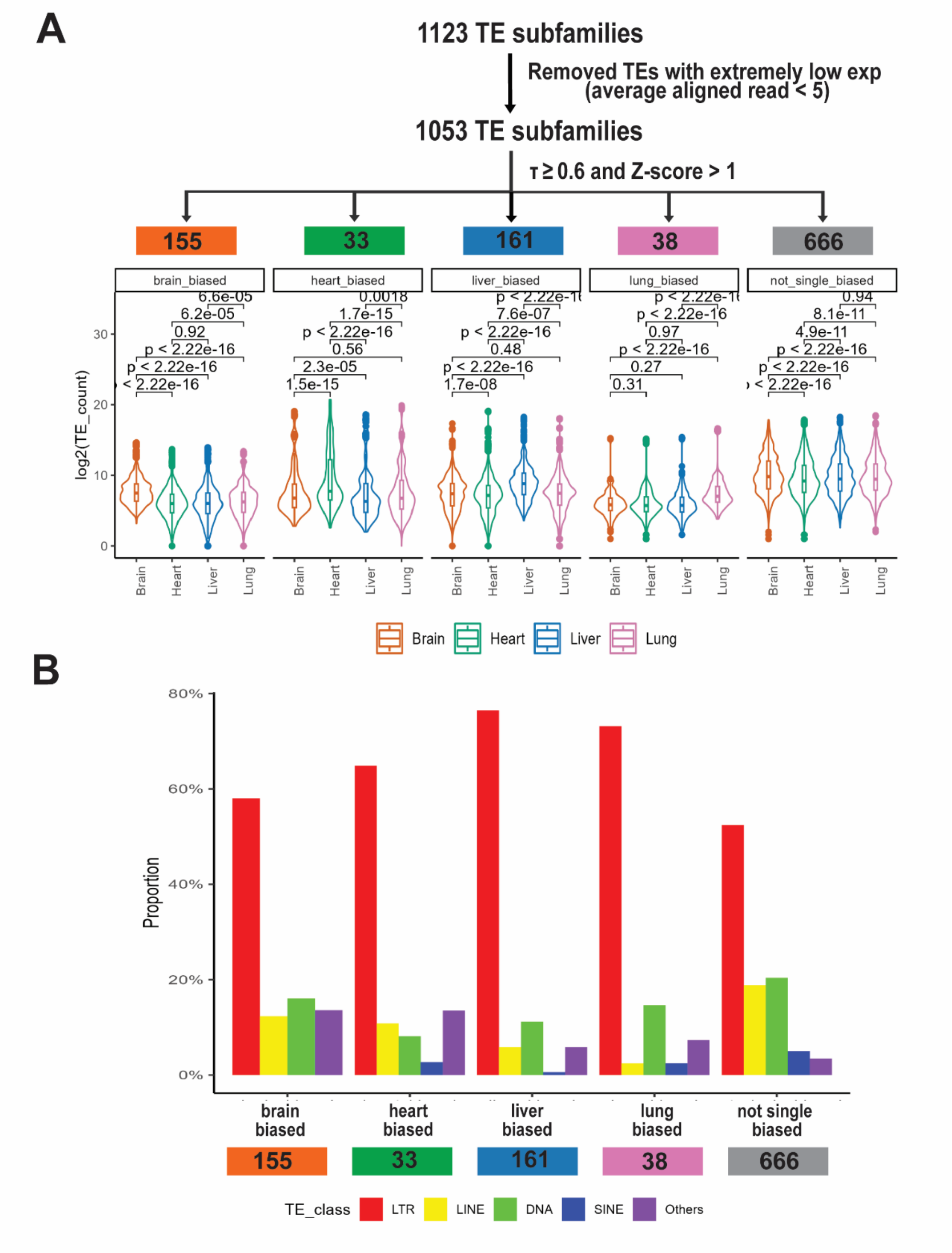


**Figure S3. Summary of TE classes of organ-biased TE subfamilies. (A)** Identification of organ-biased TE subfamilies. **(B)** Proportions of expressed TEs by class and organ shows overall dominance of LTRs and potential underrepresentation of LINE in lung and liver-biased TEs.

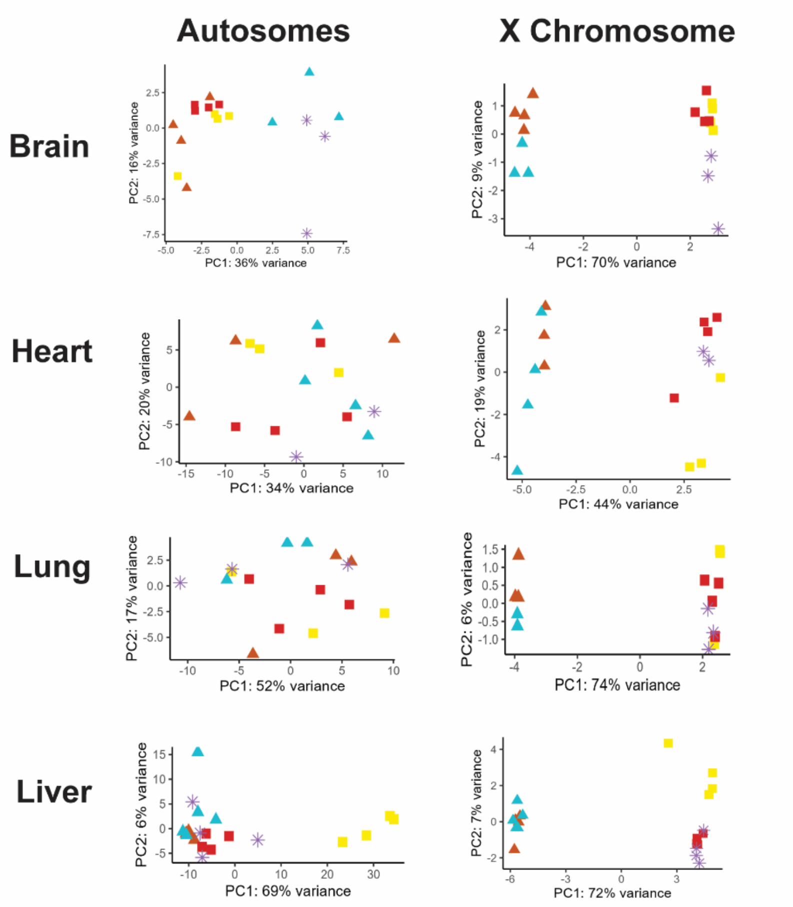


| | Brain | Heart | Lung | Liver |
|---|---|---|---|---|
| ✳ XO.F | 3 | 2 | 3 | 4 |
| ▲ XX.F | 4 | 3 | 3 | 4 |
| ▲ XXpaf.F | 3 | 4 | 3 | 5 |
| ■ XY.F | 4 | 4 | 4 | 4 |
| ■ XY.M | 4 | 3 | 3 | 4 |

**Figure S4. Principal component analyses of autosomal and X-linked gene expression in brain, heart, lung, and liver. Top:** The 300 genes with most variable expression were used. For autosomal genes, brain samples show grouping based on genetic background, while no grouping by sex or genotype is found in heart or lung. Liver samples show grouping based on gonadal sex. For X-linked genes, all four organs show grouping of samples based on the number of X chromosomes. **Bottom**: table shows the number of samples in each sex/genotype/organ group.

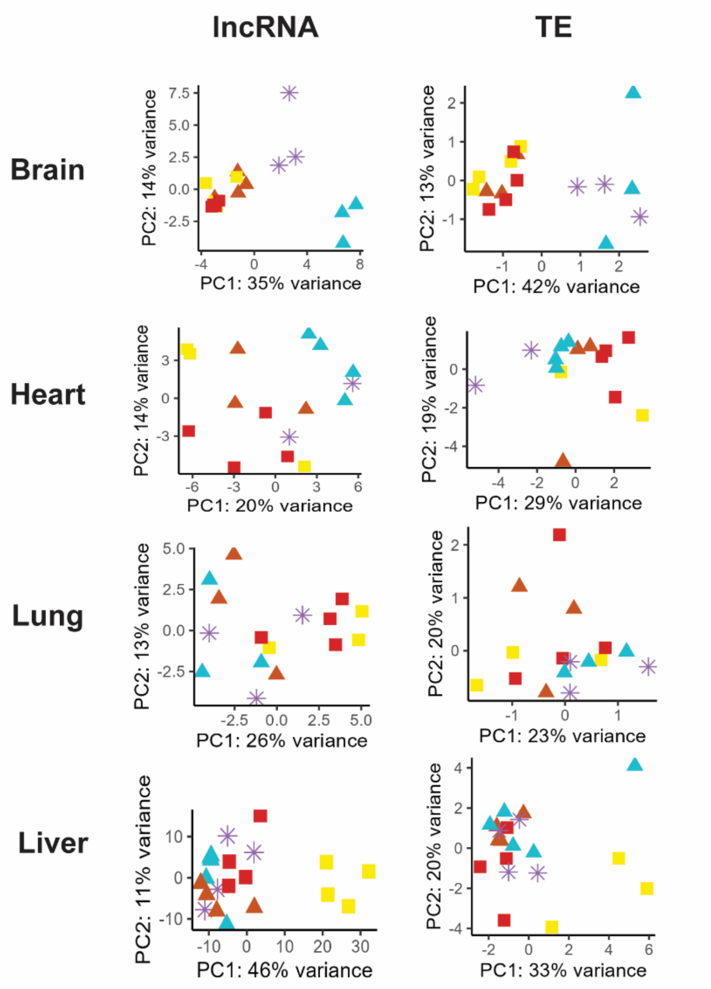


| | Brain | Heart | Lung | Liver |
|---|---|---|---|---|
| XO.F | 3 | 2 | 3 | 4 |
| XX.F | 4 | 3 | 3 | 4 |
| XXpaf.F | 3 | 4 | 3 | 5 |
| XY.F | 4 | 4 | 4 | 4 |
| XY.M | 4 | 3 | 3 | 4 |

**Figure S5. PCA plots for lncRNAs and TE instances expression in four organs. Top:** The 300 lncRNAs or TEs with most variable expression were used. Samples group by genetic background in brain, by gonadal sex in liver, and no distinct groups are observed in heart for either lncRNAs or TEs. Both autosomal and X-linked lncRNAs/TEs are used. **Bottom**: table shows the number of samples in each sex/genotype/organ group.

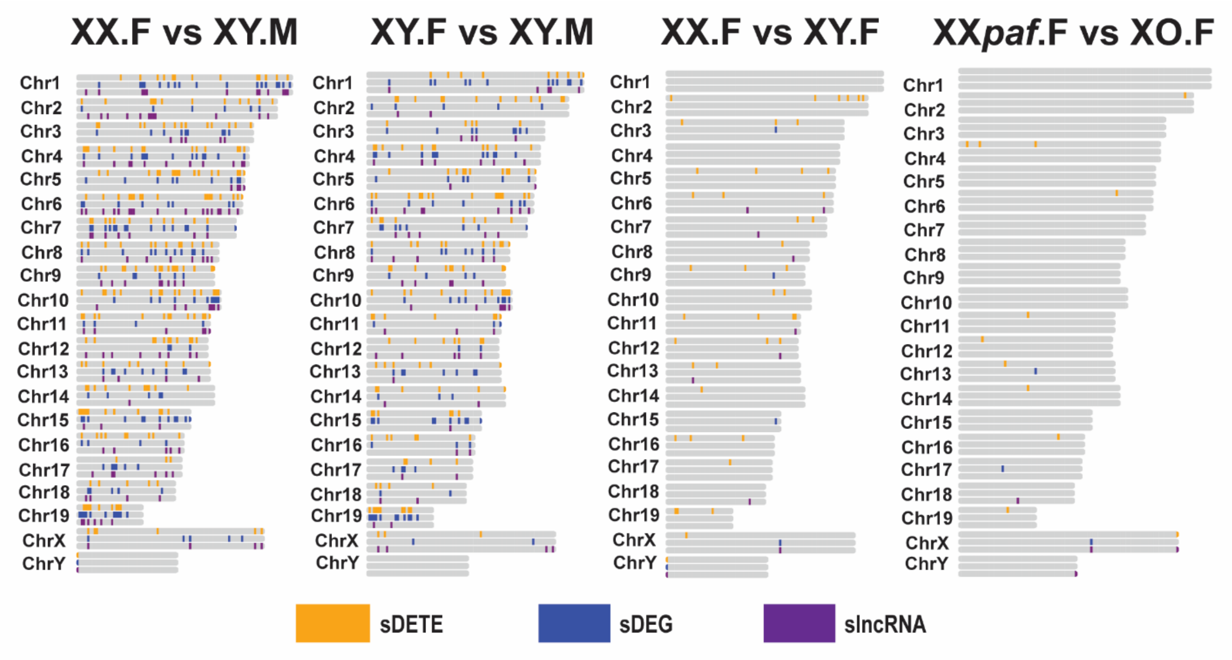


**Figure S6. Distribution of liver sDEGs, slncRNAs and sDETEs across different chromosomes in four comparisons.** Orange bars refer to location of sDETEs. Blue bars refer to locations of sDEGs. Purple bars refer to location of slncRNAs. Liver sDETEs are identified in all chromosomes (less on chrX and chrY), instead of on any specific chromosome.

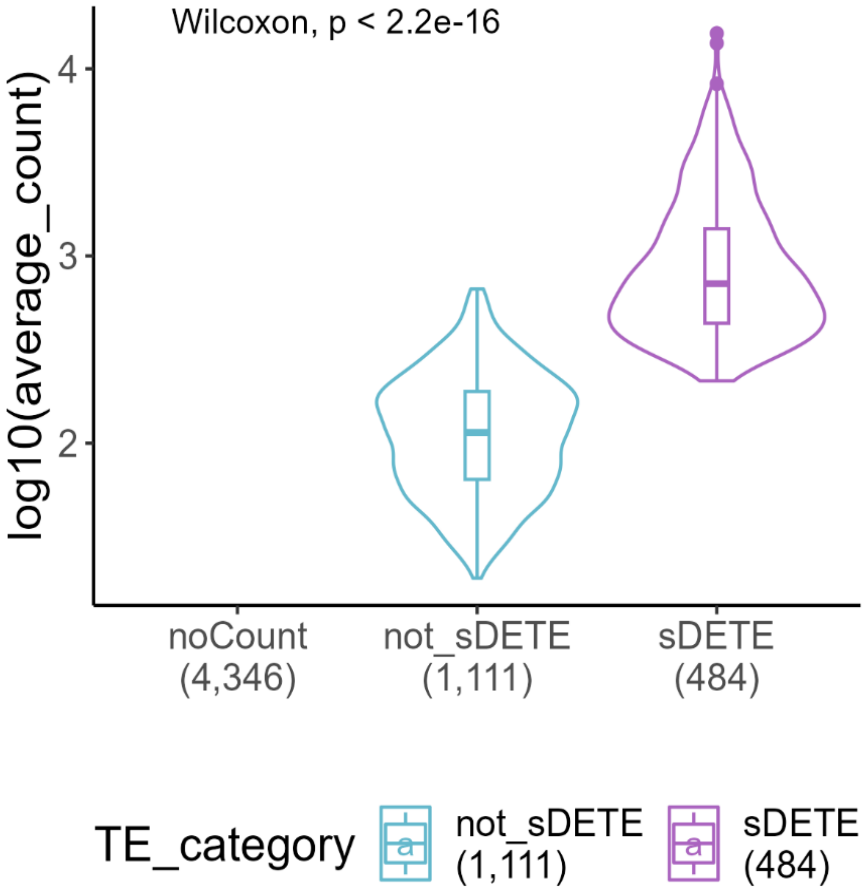


**Figure S7. Expression levels of TEs that overlap with sDEGs highlight expression variability between sDEG-overlapping TEs.** Among 5,941 sDEG-overlapping TEs, 4,346 (73%) are not expressed (no aligned reads), 1,111 (19%) are expressed but show no sex bias and 484 (8%) are sex-biased. TEs in the latter group have significantly higher expression levels than those without sex bias (Wilcoxon test).

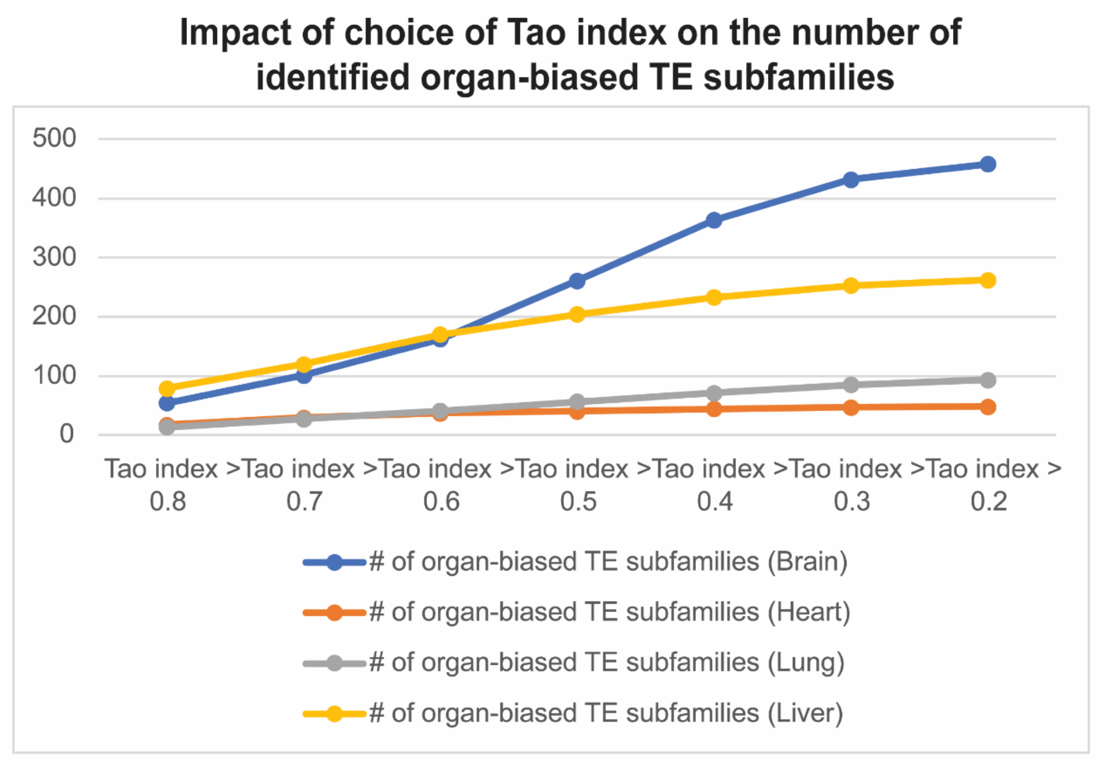


**Figure S8. Impact of choice of Tao Index on the identification of organ-biased TE subfamilies.** The cutoff threshold of z-score is set as > 1 for all organs. Each dot refers to one choice of Tao Index threshold (from Tao index $\tau > 0.8$ to $\tau > 0.2$). The Y-axis shows the number of organ-biased TE subfamilies identified. More organ-biased TE subfamilies are identified in brain and liver than in lung and heart in all settings. Brain samples start to show more organ-biased TE subfamilies than liver after the change of $\tau > 0.6$ to $\tau > 0.5$.
